# Systematic Evaluation of Signal Peptide-Driven Protein Secretion in the Fast-Growing Cyanobacterium *Synechococcus* sp. PCC 11901

**DOI:** 10.64898/2026.05.20.726548

**Authors:** José Ángel Moreno-Cabezuelo, Allanah Booth, Da Lin, Kiran Gathani, David S. Kim, Uma Shankar Sagaram

## Abstract

The fast-growing cyanobacterium *Synechococcus sp*. PCC 11901 is emerging as a promising chassis for photosynthetic biomanufacturing. Here we report recombinant protein production in PCC 11901 via signal peptide-mediated secretion, enabling direct recovery of target proteins from the culture medium without cell disruption. Seven signal peptides spanning both Sec and Tat pathways are screened using eYFP as a reporter, with secretion quantified daily over seven days by fluorescence measurements. FutA, belonging to the Tat pathway from *Synechocystis* sp. PCC 6803, achieves 92.2% extracellular export by day 7, substantially outperforming all Sec candidates, including the best Sec signal peptide thermitase from *Cyanobacterium aponinum* PCC 10605 (55.7%). Signal peptide-bearing strains exhibit growth reductions of up to 26% relative to the wild-type, with FutA most affected, indicating a general metabolic cost correlated with secretion efficiency. The best-performing signal peptides from both pathways, FutA and thermitase, are validated with secretion of lichenase. Notably, the rank order of signal peptide performance is reversed for lichenase: thermitase demonstrates 2.6-fold higher extracellular activity than FutA, indicating that optimal signal peptide selection is cargo-dependent. These results establish PCC 11901 as a secretion-competent chassis and provide a rational framework for matching signal peptide pathways to target protein properties.

## 1. Introduction

Cyanobacteria are photosynthetic prokaryotes that account for an estimated 25% of oceanic net primary production [1]. Their capacity to fix CO_2_ using solar energy, combined with minimal nutritional requirements and a rapidly expanding genetic toolkit, has positioned them as compelling platforms for sustainable biomanufacturing [2,3]. Current industrial production of recombinant proteins relies predominantly on *Escherichia coli*, which benefits from rapid growth, well-characterised genetics, and established bioprocesses at scale [4,5]. However, *E. coli*-based systems impose non-trivial costs in terms of bioreactor costs, sterility requirements, organic feedstocks, aeration, and temperature control. Cyanobacteria circumvent several of these limitations: they require only light, inorganic carbon (in the form of CO_2_ or NaHCO3 typically), nitrates, phosphates, water, and trace minerals, and have already demonstrated the capacity to produce commercially relevant proteins at laboratory scale at low cost [6–8]. Moreover, recent advances in developing genetic toolkits for fast-growing cyanobacteria have paved the way for metabolic engineering and recombinant expression to be possible in these non-model organisms [9].

Among the expanding repertoire of genetically tractable cyanobacterial strains, *Synechococcus* sp. PCC 11901 (hereafter PCC 11901), has emerged as a particularly promising production chassis. Originally isolated from the Johor river estuary in Singapore and characterised as a fast-growing, halotolerant strain [10], PCC 11901 exhibits a doubling time of approximately two hours and sustains growth to cell densities exceeding 33 g dry cell weight (DCW) L^−1^ [10,11]. The PCC 11901 genome (3.08 Mb; 3,316 predicted protein-coding genes including plasmids; GenBank CP040360.1) [10] has been the subject of comparative genomic analysis, which revealed a simplified electron transport chain and a reduced phycobilisome light-harvesting antenna relative to model cyanobacteria, features proposed to underlie its reduced susceptibility to photoinhibition and elevated photosynthetic and respiratory rates [11]. Integrated photo-mechanistic modelling subsequently confirmed that PCC 11901 achieves more than two-fold faster growth than *Synechocystis* sp. PCC 6803 under optimised illumination at approximately 735 µmol photons m^−2^ s^−1^ [12]. Most recently, comprehensive genetic engineering toolboxes have been established for PCC 11901, including validated neutral integration sites, constitutive and inducible promoter systems, CRISPRi-based gene repression, and CRISPR-Cas12a-mediated genome editing [13,14]. Taken together, these attributes make PCC 11901 exceptionally well suited for industrial-scale biotechnology applications.

Despite these advances, a fundamental bottleneck constrains the industrial deployment of cyanobacterial protein production: recombinant products overwhelmingly accumulate inside the cell [15]. Recovery of intracellular protein requires cell lysis, whether mechanical, chemical, or enzymatic, followed by multi-step chromatographic purification. Downstream processing typically accounts for 50-80% of total bioprocess costs [16,17], and obligatory cell destruction precludes the continuous harvesting strategies that would substantially improve process economics. Engineering strains to secrete target proteins directly into the culture medium would bypass this bottleneck, enabling simplified purification workflows and, potentially, continuous product recovery from growing cultures.

Protein secretion in cyanobacteria is poorly characterised relative to well-studied heterotrophic bacteria. Only four of the eleven recognised bacterial secretion systems have been identified in cyanobacteria: the Type I (T1SS), Type IV (T4SS), Type V (T5SS), and Type IV pilus assembly (T4P) systems [18]. Moreover, the outer membrane of *Synechocystis* sp. PCC 6803 exhibits permeability approximately 20-fold lower than that of *E. coli* [19], a property that may be related in part to the presence of an S-layer, a paracrystalline protein lattice that forms the outermost layer of the cell envelope, in this species [20,21]. This low permeability has contributed to the prevailing view that cyanobacteria are inherently poor secretors. However, this paradigm has recently been challenged by experimental evidence demonstrating that cyanobacteria can secrete heterologous proteins, including enzymes with complex folding requirements for functionality, at quantifiable levels. Russo et al. (2019) reported secretion of a lytic polysaccharide monooxygenase at 779 ± 40 µg L^−1^ from *S. elongatus* UTEX 2973 [22], and a highly sensitive NanoLuc-based secretion reporter subsequently enabled quantitative analysis of two-step protein secretion in *Synechocystis* sp. PCC 6803 [23]. More broadly, deep exoproteomic analysis using the EXCRETE workflow revealed that up to 85% of all potentially secreted proteins could be identified in the cyanobacterial exoproteome, with cell envelope maintenance and nutrient acquisition representing conserved core functions of the secretome across freshwater, marine, and terrestrial species [24].

Proteomic analyses have confirmed that the majority of proteins exported beyond the cytoplasmic membrane transit via two principal pathways: the general secretory pathway (Sec), which translocates unfolded proteins through the SecYEG translocon, and the twin-arginine translocation pathway (Tat), which uniquely transports fully folded proteins [18,24,25]. In both routes, an N-terminal signal peptide directs the nascent polypeptide to the appropriate translocon and is proteolytically cleaved upon arrival in the periplasm. Pioneering work by Sergeyenko *et al*. (2003) demonstrated that fusion of signal peptides to the reporter enzyme lichenase enabled secretion into the culture medium of *Synechocystis* sp. PCC 6803, providing the first proof-of-concept for heterologous protein secretion in cyanobacteria [26]. More recent studies evaluated signal peptides for heterologous secretion in *Cyanobacterium aponinum* PCC 10605, notably a cyanobacterial porin and the thermitase signal peptide [27], while Russo *et al*. (2019) demonstrated expression and secretion of a lytic polysaccharide monooxygenase in *S. elongatus* UTEX 2973 using both Sec and Tat signal peptides [22]. Subsequent work using a NanoLuc luciferase-based quantitative secretion reporter in *Synechocystis* sp. PCC 6803 demonstrated that the Type IV pilus system is not directly involved in the secretion of non-pilin proteins, indicating that the outer membrane route for two-step secretion in cyanobacteria remains to be identified [23]. A comprehensive review of cyanobacterial protein translocation systems has further highlighted the gap between the diversity of secreted proteins and the limited number of characterised secretion pathways [18,28].

Although a robust biotechnology platform has been established for PCC 11901 [10–13], no study has systematically compared the performance of Sec and Tat pathway signal peptides for heterologous protein secretion in this fast-growing strain. Here, we address this gap through a two-phase experimental programme: (i) screening seven signal peptides, five Sec and two Tat, for secretion of eYFP; (ii) validating the best-performing signal peptides with an industrially relevant enzyme lichenase. Lichenase (1,3-1,4-β-glucanase, EC 3.2.1.73) from *Clostridium thermocellum* was selected as the validation cargo for three reasons: it is a thermostable monomeric enzyme (∼36 kDa) that does not require cofactors or post-translational modifications for activity, providing a secretion readout independent of fluorescence; it has established industrial applications in brewing and animal feed for β-glucan degradation; and it was used as the secretion reporter in the research of Sergeyenko *et al*. (2003) in *Synechocystis* sp. PCC 6803 [26], enabling a direct cross-chassis comparison. We show that the Tat-pathway signal peptide FutA from *Synechocystis* sp. PCC 6803 substantially outperforms all Sec candidates for eYFP secretion, that the platform is transferable to an enzymatic cargo protein, and that optimal signal peptide selection is cargo-dependent..

## 2. Materials and Methods

### 2.1. Bacterial Strains and Culture Conditions

*Escherichia coli* DH5α was used for routine cloning, plasmid maintenance, and propagation of all constructs. For biparental conjugation into PCC 11901, the diaminopimelic acid (DAP)-auxotrophic donor strain *E. coli* MFDpir [29](Addgene #187385) (a generous gift from Antonio Lamb, ETH Zurich) was used as described in Section 2.3. *Synechococcus* sp. PCC 11901 was obtained from the Pasteur Culture Collection (PCC) and cultivated at 30°C in MAD medium [10] containing 1.8% NaCl under continuous illumination (200 µmol photons m^−2^ s^−1^) with orbital shaking (120 rpm). A wild-type (WT) strain and a non-secreting control (harbouring the P*cpc560* promoter driving eYFP without a signal peptide) were maintained under identical conditions. For selection of cyanobacterial transconjugants, spectinomycin was supplemented at 25 µg mL^−1^. WT cultures were grown without antibiotic selection.

Liquid starter cultures used for growth experiments and genetic manipulation were grown in 40 mL volumes in 100 mL conical flasks at 30 °C under continuous warm white LED illumination (200 µmol photons m^−2^ s^−1^) with orbital shaking at 130 rpm. Cultures were incubated in custom-built temperature-controlled growth chambers based on modified refrigeration units. These chambers were retrofitted with microcontroller-based systems to regulate CO_2_ concentration, humidity, and illumination. LED arrays were manually adjusted using a calibrated light probe, and a diffusion panel ensured uniform light distribution across the culture surface. CO_2_ levels were maintained at 5% (50,000 ppm) using a sensor-controlled feedback loop (± 5%), while humidity was controlled at approximately 80% using an integrated humidification system (± 20%). The refrigeration system enabled efficient heat dissipation generated by the high-intensity LED illumination.

### 2.2. Signal Peptide Selection and Construct Design

Seven signal peptides were selected to provide systematic coverage of the two principal cyanobacterial export pathways (Table 1). Putative signal peptides from PCC 11901 were identified by mining the PCC 11901 proteome (GenBank CP040360.1) and cross-referencing with published exoproteomic datasets [24,27]. Predicted subcellular localisation and signal peptide type (Sec/SPI, Sec/SPII, or Tat/SPI) were determined using SignalP v6.0[30], and cleavage sites were annotated accordingly. For the Sec pathway, four native PCC 11901 signal peptides were selected on the basis of their predicted extracytoplasmic localisation and functional annotations suggestive of secretion or envelope association: (1) Type IV pilin-like G/H family protein (a pilin-like protein expected to engage the T4P-associated export machinery, T4P-like hereafter), (2) Pentapeptide repeat-containing protein (predicted periplasmic localization, Pentapeptide hereafter), (3) Spore coat U domain-containing protein (predicted cell envelope association, Spore coat protein hereafter), and (4) DUF11 domain-containing protein (predicted outer membrane localization, DUF11 hereafter). In addition, one heterologous Sec signal peptide was also included: (5) the thermitase signal peptide from *C. aponinum* PCC 10605, which has been shown to drive GFP secretion in *C. aponinum* using RSF1010-based plasmids [21]. For the Tat pathway, two signal peptides carrying canonical twin-arginine motifs were selected: (6) FutA from *Synechocystis* sp. PCC 6803, a well-characterised ferric iron-binding periplasmic protein [31], and (7) PhoX, a PhoX family alkaline phosphatase native to PCC 11901.

**Table 1.** Signal peptides evaluated in this study. Pathway assignments and signal peptide types were predicted using SignalP v6.0. Seven signal peptides were selected to provide systematic coverage of the two principal cyanobacterial export pathways (Table 1; full amino acid and DNA sequences provided in **Supplementary Table S1**).

| # | Signal Peptide | Strain Origin | Type of Secretion | NCBI Reference ID |
| --- | --- | --- | --- | --- |
| 1 | Type IV pilin-like G/H family protein | *Synechococcus* sp. PCC 11901 | Sec | WP_138071863.1 |
| 2 | Pentapeptide repeat-containing protein | *Synechococcus* sp. PCC 11901 | Sec | WP_138071668.1 |
| 3 | PhoX family protein | *Synechococcus* sp. PCC 11901 | Tat | WP_138072907.1 |
| 4 | Spore coat protein U domain-containing protein | *Synechococcus* sp. PCC 11901 | Sec | WP_138071630.1 |
| 5 | DUF11 domain-containing protein | *Synechococcus* sp. PCC 11901 | Sec | WP_138073633.1 |
| 6 | Thermitase | *Cyanobacterium aponinum* PCC 10605 | Sec | WP_099435530.1 |
| 7 | FutA Iron uptake protein A2 | *Synechocystis* sp. PCC 6803 | Tat | BAA10591.1 |

All coding sequences were codon-optimised for the PCC 11901 codon usage table (derived from the GenBank CP040360.1 proteome) and synthesised by Twist Bioscience. Signal peptide–target gene fusions were assembled in-frame by Golden Gate assembly into the self-replicating broad-host-range plasmid RSF1010. The RSF1010-based expression system was selected over chromosomal integration at the *mrr* neutral site based on preliminary experiments in our laboratory indicating that self-replicating plasmids yielded higher and more consistent recombinant protein expression in PCC 11901 (data presented in the discussion, section 4). The general construct architecture was: P*cpc560* promoter – signal peptide – target gene – native terminator from *E. coli* ECK120010850. Upon secretion into the periplasm, the signal peptide is cleaved, leaving only the mature target protein. Correct assembly was verified by full plasmid sequencing.

### 2.3. Biparental Conjugation

RSF1010-derived constructs were introduced into PCC 11901 by biparental conjugation using the diaminopimelic acid (DAP)-auxotrophic donor strain *E. coli* MFDpir (Addgene #187385). The cargo plasmid was transformed into chemically competent MFDpir cells by heat shock and selected on LB agar supplemented with 0.3 mM DAP and spectinomycin (25 µg mL^−1^). For conjugation, overnight cultures of the transformed donor were washed three times with LB + DAP without antibiotics. PCC 11901 cultures (OD_730_ 0.5–2.0) were washed four times with fresh MAD medium. Equal volumes (900 µL) of donor and recipient were combined, mixed gently, and incubated at room temperature for 4–5 h. The mating mixture was pelleted (1,500 × *g*, 10 min), resuspended in residual supernatant, and was spread onto 0.45 µm membrane filters (Millipore HATF) placed on LB:MAD (1:1) agar containing DAP without antibiotics. After 24 h under illumination, membranes were transferred to MAD agar containing spectinomycin (25 µg mL^−1^) and lacking DAP, thereby counter-selecting the donor. Conjugant colonies appeared within 5–7 days and were verified by plasmid sequencing and sustained growth on selective medium.

### 2.4. Culture Conditions for Secretion Assays

Each transconjugant strain, together with the non-secreting control, was inoculated into MAD liquid medium (1.8% NaCl, 25 µg mL^−1^ spectinomycin). The WT control was cultured in identical medium without spectinomycin. All strains were cultured for 7 days at 30 °C in a CO_2_-controlled incubator maintained at 5% CO_2_ (v/v) under continuous illumination at (200 µmol photons m^−2^ s^−1^) with shaking at (130 rpm). n = 3 biological replicates. Samples were collected daily from day 1 through day 7 for subcellular fractionation and fluorescence quantification. Growth of all transconjugant strains was monitored by OD_730_ measurements at each sampling point to assess potential fitness costs associated with signal peptide expression.

### 2.5. Subcellular Fractionation

To measure the intracellular and secreted protein fractions, aliquots of 2 mL from day 1 to day 7 cultures were collected in duplicate and OD_730_ was recorded at each time point. Cells were pelleted by centrifugation (9,500 × *g*, 10 min, 4 °C). The supernatant was retained as the extracellular fraction. The cell pellet was washed 5 times with 2 mL fresh MAD medium (10 min per wash) to remove residual extracellular protein and then resuspended in 1 mL MAD medium. Fluorescence of both supernatant and cell fractions was measured as described in Section 2.6.

### 2.6. Fluorescence Quantification

For each fraction, 200 µL aliquots were dispensed in triplicate into 96-well Nunc™ plates. Fluorescence intensity was measured on a Tecan Infinite 200 PRO (Tecan Group Ltd., Männedorf, Switzerland) microplate reader using bottom-reading mode (excitation: 488 nm, bandwidth 9 nm; emission: 530 nm, bandwidth 20 nm; 25 flashes per well; gain: optimal; multiple reads per well in a 2 × 2 square-filled pattern). The excitation wavelength of 488 nm was selected rather than the eYFP excitation maximum (513 nm) to maintain sufficient spectral separation between excitation and emission wavelengths, as recommended for monochromator-based plate readers when excitation and emission peaks are closely spaced. Background fluorescence from WT fractions was subtracted from all readings. Fluorescence values were then divided by OD_730_ to yield relative fluorescence units per optical density (RFU/OD).

### 2.7. Calculation of Secretion Efficiency

The percentage of protein secreted to the extracellular space was calculated as:

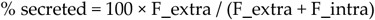

where F_extra denotes the background-corrected fluorescence of the cell-free supernatant (extracellular fraction) and F_intra denotes the background-corrected fluorescence of the washed, resuspended cell pellet (intracellular fraction), both normalised to OD_730_.

### 2.8. Qualitative Imaging

Extracellular fractions, and intracellular fractions were photographed under blue-light (488 nm) excitation using a *ChemiDoc* imaging system (*Bio-Rad*) equipped with the appropriate emission filter for eYFP.

### 2.9. Lichenase Secretion and Activity Assays

The lichenase gene (*licB*) from *Clostridium thermocellum*, encoding a 1,3–1,4-β-glucanase, was fused in-frame with the FutA and thermitase signal peptides using the same RSF1010 construct architecture as in Phase 1 (P*cpc560* – signal peptide– *licB* – terminator). An additional construct expressing lichenase fused to its native *C. thermocellum* signal peptide was included as a dual-purpose control: this Gram-positive signal peptide was not expected to be recognised by the cyanobacterial secretion machinery, thereby serving simultaneously as a positive control for intracellular lichenase expression and a negative control for extracellular secretion. The wild-type strain (lacking the *licB* gene) served as an additional negative control. Constructs were assembled by Golden Gate assembly, verified by full plasmid sequencing, and introduced into PCC 11901 by biparental conjugation as described in Section 2.3. Expression and culture conditions were identical to Phase 1 (Section 2.4).

Qualitative assessment of extracellular lichenase activity was performed using two complementary Congo Red plate assay formats adapted from Sergeyenko *et al*.[26]. In the colony overlay format, transconjugant cells were grown on MAD agar supplemented with 25 µg mL^−1^ spectinomycin for 3 days at 30°C under continuous illumination. Colonies were overlaid with top agar (0.7% agarose in 50 mM Tris-HCl pH 8.0) supplemented with 0.05% lichenan (*Sigma-Aldrich, G6513*). After polymerisation, plates were incubated at 65°C for 4–6 h to allow the thermostable lichenase to digest the substrate while inactivating endogenous mesophilic enzymes. Plates were then stained with 0.5% Congo Red for 10 min at room temperature with gentle shaking and destained three times with 1 M NaCl for 5 min each. Lichenase activity was visualised as clear zones against a dark red background of undigested lichenan. In the liquid culture format, cell-free supernatants were prepared as described in Section 2.5. Filtered supernatants were mixed with an equal volume of MAD medium containing 3% agar, poured into Petri dishes and allowed to solidify. Plates were then overlaid with lichenan-containing top agar and processed as above.

Quantitative assessment of extracellular lichenase activity was performed using the *Megazyme* Malt β-Glucanase/Lichenase Assay Kit (K-MBG4, *Megazyme*, Bray, Ireland) following the manual lichenase assay procedure. The kit employs the chromogenic substrate BCNPBG4 (4,6-*O*-benzylidene-2-chloro-4-nitrophenyl-β-(3^1^-β-D-cellotriosyl-glucoside)), which is specifically cleaved by lichenase (EC 3.2.1.73) to release 2-chloro-4-nitrophenol (CNP). The benzylidene acetal blocking group prevents hydrolysis by exo-acting enzymes including β-glucosidase and cellobiohydrolase, ensuring assay specificity. Cell-free supernatants were prepared as described in Section 2.5 and assayed undiluted. MBG4 substrate (100 µL) was dispensed into 13 mL glass tubes and pre-incubated at 40°C for 3 min. Sample (200 µL) was added, vortexed, and incubated at 40°C for exactly 10 min. The reaction was terminated by addition of 3,000 µL stopping reagent (2% w/v Tris buffer, pH 10.0), which simultaneously develops the yellow phenolate colour of released CNP. Absorbance was measured at 400 nm against distilled water. For each strain, a sample blank was prepared by adding the stopping reagent prior to the sample and substrate (to correct for background absorbance of the culture supernatant), and a reagent blank was prepared using Buffer D (100 mM sodium phosphate, pH 6.5) in place of the sample. The assay was validated using a *Bacillus* sp. lichenase standard supplied with the kit (recovery: 92%). Lichenase activity was calculated as MBG4 U/mL = ΔE_400_× 0.0994 x 50, where ΔE_400_= Abs(reaction) − Abs(sample blank) − Abs(reagent blank), where 50 is the Megazyme conversion factor relating activity to the original enzyme preparation and one MBG4 Unit is defined as the amount of enzyme releasing one micromole of CNP per minute (εmM = 16.6 at 400 nm in 2% Tris buffer, pH 10.0). Activity values were normalised to the optical density of the culture at harvest (OD_730_) and expressed as mU/mL/OD_730_.

### 2.10. Use of Generative Artificial Intelligence

During the preparation of this manuscript, no generative artificial intelligence (GenAI) tools were used for data analysis, experimental design, or interpretation of results. AI-assisted writing tools were used for language editing and manuscript preparation. The authors reviewed and take full responsibility for the content of this publication.

### 2.11. Statistical Analysis

All experiments were performed with a minimum of three biological replicates. Data is presented as mean ± (SD). Statistical analyses were performed using GraphPad Prison v.11.0.0 and are detailed alongside figures.

## 3. Results

### 3.1. Phase 1: A Systematic Signal Peptide Screen Identifies FutA and Thermitase as Optimal Leaders for eYFP Secretion

To evaluate the secretion capacity of PCC 11901, seven signal peptides, five targeting the Sec pathway (thermitase, T4P-like, DUF11, spore coat protein, pentapeptide) and two targeting the Tat pathway (FutA, PhoX), were fused to the N-terminus of eYFP and expressed from the self-replicating plasmid RSF1010 under the constitutive P*cpc560* promoter. The architecture of the seven resulting constructs, showing the shared backbone elements together with the signal peptide-specific expression cassettes (P*cpc560* – signal peptide –eYFP – terminator), is presented in **Figure 1A**. Cultures were sampled daily from day 1 to day 7, and eYFP fluorescence was quantified in both the extracellular and intracellular fractions at each time point. Growth of all transconjugant strains was monitored in parallel.

**Figure 1.**
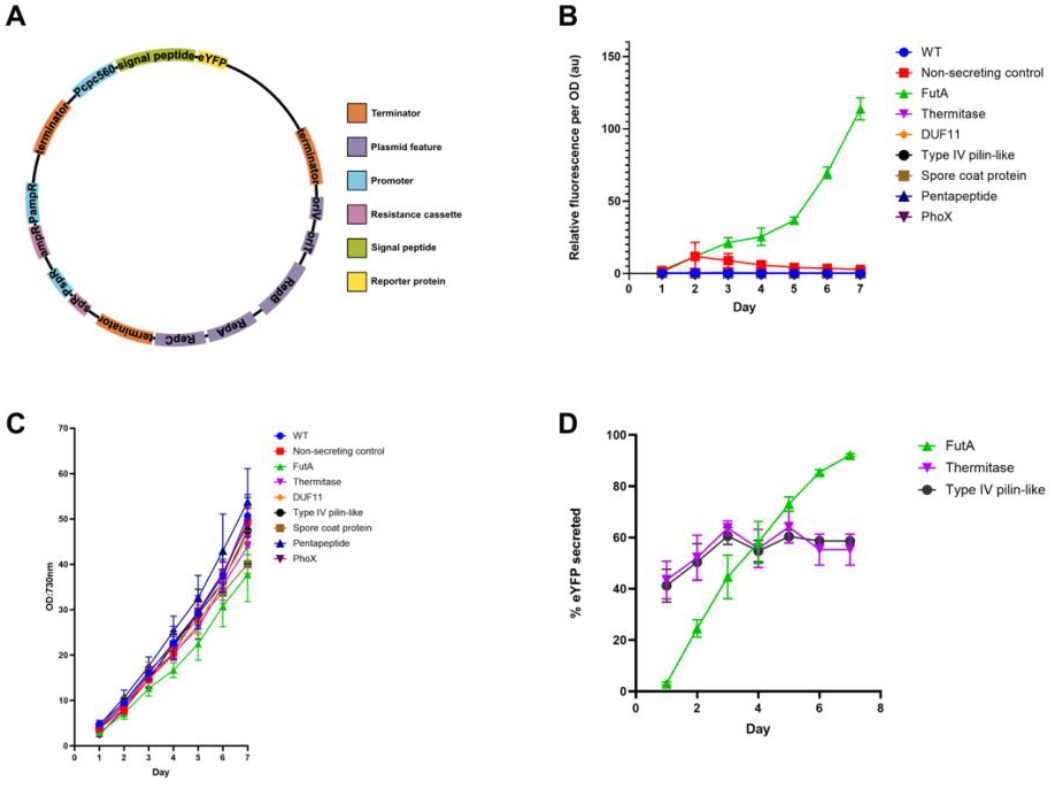
Systematic signal peptide screen for eYFP secretion in *Synechococcus* sp. PCC 11901. **(A)** Architecture of the RSF1010-based expression construct; signal peptide variants are detailed in Table 1. **(B)** Extracellular eYFP fluorescence normalised to OD_730_ from day 1 to day 7. **(C)** Growth curves (OD_730_) over the 7-day time course. **(D)** Time-resolved secretion efficiency (% eYFP secreted) for the three highest-performing signal peptides: FutA (Tat), thermitase (Sec), and T4P-like (Sec). Error bars = SD, n = 3 biological replicates. Made in GraphPad Prism and Affinity Designer 2.

All seven signal peptide constructs secreted detectable eYFP fluorescence to the extracellular space over the 7-day time course (**Figure 1B**), demonstrating that PCC 11901 is competent for recombinant protein export via both the Sec and Tat translocation routes. The wild-type strain did not exhibit extracellular fluorescence above background at any time point. The non-secreting control showed low-level extracellular fluorescence (9.3 ± 2.9% at day 7), attributable to basal cell lysis. Signal peptide constructs vastly exceeded this baseline, FutA reached 113.8 RFU/OD at day 7, approximately 40-fold higher and the signal peptide-dependent rank order is inconsistent with a lysis-based mechanism. Furthermore, the outer membrane of cyanobacteria lacks classical porins and exhibits >20-fold lower permeability to small organic solutes than *E. coli* [19]. eYFP is a folded ∼27 kDa protein and so passive leakage is highly implausible. Intracellular fluorescence measurements confirmed that all signal peptide-bearing strains maintained eYFP production throughout the time course (**Supplementary Figure S1**), while whole-culture fluorescence is shown in **Supplementary Figure S2**. Progressive accumulation of secreted eYFP by FutA is also visible in *ChemiDoc* UV images of the extracellular fraction from day 1 to day 7 (**Supplementary Figure S3**).

The secretion efficiency of each signal peptide, defined as the percentage of total eYFP fluorescence detected in the extracellular fraction, was determined at day 7 from three biological replicates (**Supplementary Figure S4)**. Among the Sec-pathway candidates, the T4P-like signal peptide native to PCC 11901 achieved the highest efficiency (58.7 ± 0.1%), closely followed by the heterologous thermitase signal peptide from *C. aponinum* PCC 10605 (55.7 ± 3.5%). The remaining native PCC 11901 Sec signal peptides yielded lower efficiencies: pentapeptide (20.1 ± 1.4%), DUF11 (14.3 ± 0.7%), and spore coat protein (11.0 ± 1.6%). Notably, DUF11 directed only 14.3% of eYFP to the extracellular fraction despite its predicted outer membrane localisation.

Strikingly, the Tat-pathway candidate FutA from *Synechocystis* sp. PCC 6803 achieved the highest secretion efficiency of any signal peptide tested, reaching 92.2 ± 0.3% extracellular export by day 7, substantially exceeding all Sec candidates and the second Tat candidate PhoX (17.1 ± 0.3%)). Extracellular fluorescence at day 7 for all strains is shown in **Supplementary Figure S5**. These results identified FutA as the leading signal peptide overall. Of the two highest efficiency Sec candidates, the thermitase tag showed a higher amount of eYFP secreted (0.26 ± 0.053 RFU/OD) than the T4P-like tag (0.23 ± 0.026 RFU/OD). Notably, the superior performance of FutA, a cyanobacterial signal peptide, compared to heterologous alternatives suggests that non-native signal peptides may be better adapted to the host secretion machinery in cyanobacterial systems. FutA and thermitase were thus both carried forward for validation with an additional target protein.

### 3.2. Secretion Kinetics Reveal Distinct Temporal Profiles for Tat and Sec Leaders

FutA-driven extracellular fluorescence increased progressively from 0.93 RFU/OD at day 1 to 113.8 RFU/OD at day 7 **(Figure 1B)**, with secretion efficiency reaching 92.2% without saturation (**Figure 1D**). By contrast, thermitase and T4P-like reached near-plateau efficiencies by day 3 (∼64% and ∼61% respectively), declining slightly to 55.7% and 58.7% by day 7 (**Figure 1D**). PhoX demonstrated 17.1 ± 0.3% efficiency by day 7, further suggesting that PCC 11901 can export heterologous proteins via the Tat pathway but that signal peptide identity, rather than pathway assignment alone, is the primary determinant of secretion efficiency. Complementary visualisations as grouped bars per day are provided for extracellular fluorescence (**Supplementary Figure S6**) and secretion efficiency (**Supplementary Figure S7**).

### 3.3. Signal Peptide Expression Imposes a Growth Cost Correlated with Secretion Efficiency

All signal peptide-bearing transconjugant strains exhibited a growth reduction relative to the wild-type and non-secreting controls over the 7-day culture period (**Figure 1C**). At day 7, OD_730_ values were 50.7 ± 1.2 for the wild-type and 49.4 ± 1.0 for the non-secreting control, compared with 37.7 ± 1.4 for FutA, 43.9 ± 1.2 for thermitase, 47.4 ± 1.6 for T4P-like, and 48.8 ± 1.1 for PhoX. The growth reduction was most pronounced for FutA, which reached a final OD_730_ approximately 26% lower than the wild-type, consistent with a substantial metabolic cost associated with high-level protein secretion through the Tat pathway.

An inverse correlation was observed between secretion efficiency and final cell density: the three highest-secreting strains, FutA (92.2%; OD_730_ = 37.7), T4P-like (58.7%; OD_730_ = 47.4), and thermitase (55.7%; OD_730_ = 43.9), reached lower final densities than the lowest-secreting strains, pentapeptide (20.1%; OD_730_ = 53.8) and spore coat (11.0%; OD_730_ = 40.1). This growth penalty was observed across both Sec and Tat pathway constructs, suggesting that the metabolic burden arises from engagement of the translocation machinery and diversion of cellular resources towards protein export, rather than from pathway-specific toxicity.

The non-secreting control accumulated the highest intracellular eYFP fluorescence of any strain (27.3 RFU/OD at day 7) and growth was comparable to the wild-type (OD_730_ = 49.4 vs 50.7), indicating that high-level eYFP expression *per se* does not impose a significant growth cost. Rather, translocation across the membrane seems to incur the observed fitness penalty. The intracellular fluorescence in the non-secreting control declined from a peak of 65.6 RFU/OD at day 2 to 27.3 RFU/OD at day 7. As the P*cpc560* promoter is constitutive and not density-dependent, this decline is unlikely to result solely from growth dilution and may reflect reduced promoter activity under high cell-density conditions, intracellular protein turnover, or increased metabolic burden at later growth stages.

### 3.4. Tat vs Sec Pathway Comparison: FutA Outperforms All Sec Leaders for eYFP

The inclusion of signal peptides targeting both major translocation pathways enabled a direct pathway comparison. The Tat pathway translocates fully folded proteins across the cytoplasmic membrane, permitting cytoplasmic chromophore maturation in eYFP prior to translocation, thereby circumventing the challenges of fluorescent protein folding in the oxidising periplasmic environment [32,33]. FutA (Tat) yielded 92.2 ± 0.3% secretion efficiency at day 7, compared with 58.7 ± 0.1% for T4P-like (Sec) and 55.7 ± 3.5% for thermitase (Sec), a difference that was statistically significant by one-way ANOVA with Tukey’s multiple comparisons test (F(7, 16) = 282.3, p < 0.0001; **Supplementary Figure S4)**. This pattern was also reflected in absolute extracellular fluorescence levels (**Supplementary Figure S5**; one-way ANOVA with Tukey’s multiple comparisons test, F(8, 18) = 630.1, p < 0.0001). This advantage was consistent across all time points from day 3 onwards, and the gap between FutA and the best Sec candidates widened progressively, reaching 34–37 percentage points by day 7.

Our experimental design included five Sec and two Tat signal peptides. This asymmetry means that while FutA clearly outperforms all individual Sec candidates tested, the present data do not permit a generalised conclusion about intrinsic Tat pathway superiority.

### 3.5. Phase 2: Lichenase Secretion Validates Enzymatic Protein Export

Having identified FutA and thermitase as optimal signal peptides, for Tat and Sec pathways, respectively, we tested whether the platform could deliver enzymatically active proteins to the extracellular space. Lichenase (1,3–1,4-β-glucanase) from *Clostridium thermocellum*, an enzyme of industrial relevance in brewing and animal feed for β-glucan degradation, was selected because it provides an activity-based secretion readout independent of fluorescence, and because it was used as the secretion reporter in the seminal study by Sergeyenko *et al*. (2003) in *Synechocystis* sp. PCC 6803 [26], enabling a direct cross-chassis comparison. In that study, lichenase fused to positively charged pilin-type leader peptides (*PilA1* and *Slr2016*) was secreted from chromosomally integrated cassettes, whereas lichenase expressed without a leader peptide remained intracellular, as did lichenase fused to an artificial leader carrying an overall negative charge (−4 at pH 7) [26].

Lichenase constructs were generated with both FutA (Tat) and thermitase (Sec) signal peptides and expressed from the RSF1010 plasmid under the P*cpc560* promoter. As a dual-purpose control, we included a construct expressing lichenase fused to its native *C. thermocellum* signal peptide. This signal peptide, which evolved in a Gram-positive thermophilic bacterium, was not expected to be recognised by the cyanobacterial secretion machinery and therefore served simultaneously as a positive control for intracellular lichenase expression and as a negative control for extracellular secretion. The wild-type strain (lacking the *licB* gene) served as an additional negative control.

Extracellular lichenase activity was first assessed qualitatively using Congo Red plate assays adapted from Sergeyenko *et al*. (2003) [26]. In the colony overlay format, clear zones surrounded FutA and thermitase colonies, while the native *C. thermocellum* signal peptide construct showed clearing only beneath the colony, consistent with intracellular expression without extracellular export (**Supplementary Figure S8**). In the liquid culture format, cell-free supernatants from FutA-lichenase and thermitase-lichenase strains showed clear zones indicating extracellular lichenase activity, while wild-type and native *C. thermocellum* signal peptide control showed no extracellular activity (Figure 2A). Further, quantitative assessment using the *Megazyme* K-MBG4 chromogenic assay confirmed these findings (**Figure 2B**; raw absorbance data at 400 nm shown in **Supplementary Figure S9**). When normalised to culture density, thermitase-lichenase exhibited a mean extracellular activity of approximately 74.99 ± 1.78 mU/mL/OD_730_, while FutA-lichenase showed a mean activity of approximately 28.83 ± 0.96 mU/mL/OD_730_. Both the wild-type and the native signal peptide control showed no detectable extracellular activity (ΔE_400_ ≤ 0 after blank subtraction). Differences between strains were highly significant by one-way ANOVA with Tukey’s multiple comparisons test (F(3, 8) = 299.2, p < 0.0001).

**Figure 2.**
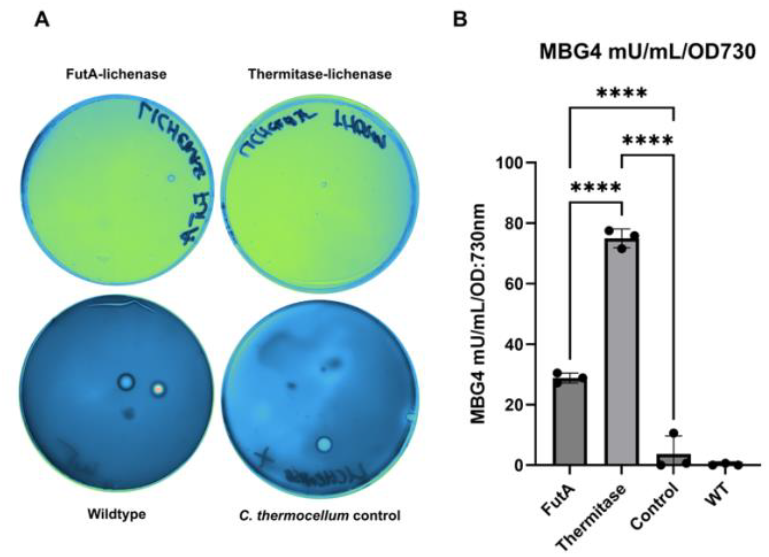
Lichenase secretion in *Synechococcus* sp. PCC 11901. **(A)** Liquid culture Congo Red assay: cell-free supernatants embedded in agar from FutA-lichenase (top left) and thermitase-lichenase (top right) show clear zones after Congo Red staining, indicating extracellular lichenase activity. Wild-type (bottom left) and native *C. thermocellum* signal peptide control (bottom right) show no extracellular activity. **(B)** Quantitative extracellular lichenase activity (mU/mL/OD_730_) measured by the *Megazyme* K-MBG4 assay. Data were analysed using oa one-way ANOVA with Tukey’s multiple comparisons test (F(3,8) = 299.2, **** = p<0.0001). Error bars = SD, n = 3 biological replicates. Made in GraphPad Prism and Affinity Designer 2.

Notably, the rank order of signal peptide performance was reversed relative to Phase 1: whereas FutA substantially outperformed thermitase for eYFP secretion (92.2% vs 55.7%), thermitase directed 2.6-fold higher extracellular lichenase activity than FutA for lichenase. This cargo-dependent reversal suggests that the efficiency of a given signal peptide depends on the biophysical properties of the target protein. Lichenase from *C. thermocellum* is a monomeric enzyme that does not require cofactors or post-translational modifications for activity and may therefore be efficiently translocated in an unfolded state through the Sec pathway. By contrast, eYFP requires cytoplasmic folding and chromophore maturation to become fluorescent, properties that favour the Tat pathway, which transports fully folded substrates.

## 4. Discussion

This study establishes *Synechococcus* sp. PCC 11901 as a secretion-competent platform for extracellular recombinant protein production under industrially relevant CO_2_-supplemented growth conditions (5%) and identifies the Tat-pathway signal peptide FutA as the most effective leader for heterologous protein export in this strain. By combining a systematic signal peptide screen with daily secretion profiling and validation across two functionally distinct target proteins, we define a rational framework for engineering protein secretion in a fast-growing cyanobacterium.

As noted in the introduction, the traditional view that cyanobacteria are inherently poor secretors has already been challenged by recent exoproteomic data [24] and was based partly on the low outer membrane permeability reported for *Synechocystis* sp. PCC 6803 [19]. Our results provide direct experimental support for this revised perspective in PCC 11901. An extracellular secretion efficiency of 92.2% for FutA-driven eYFP export exceeds the qualitative secretion observed with pilin-type leaders in PCC 6803 [26] and the secretion yields reported for a lytic polysaccharide monooxygenase (LPMO) in *S. elongatus* UTEX 2973 [22], indicating that PCC 11901 possesses a secretory capacity that is among the highest reported for cyanobacteria. This finding aligns with recent deep exoproteomic analyses that identified FutA2 as one of the most abundant secreted proteins in *Synechococcus* sp. PCC 11901 and revealed that the cyanobacterial secretome is far more extensive than previously recognized [24]. We note that the FutA signal peptide used here derives from the *Synechocystis* sp. PCC 6803 FutA2 protein (characterised as a ferric iron-binding protein by Badarau et al. [31]), rather than a native PCC 11901 orthologue; nonetheless, the high secretion efficiency observed suggests strong functional compatibility with the PCC 11901 Tat translocon. The closest orthologue of FutA2 in PCC 11901 is a Fe(3+) ABC transporter substrate-binding protein (WP_138071506.1; 50.9% amino acid identity), which carries a Sec-type rather than a Tat-type signal peptide, lacking the canonical twin-arginine motif. PCC 11901 therefore does not possess a native Tat-dependent iron uptake signal peptide, and a heterologous leader from PCC 6803 was required to evaluate Tat-mediated secretion in this strain. What might account for the permissive secretion phenotype of PCC 11901? Comparative genomics revealed a streamlined electron transport chain relative to PCC 6803 [11], and differences in cell envelope-related gene content have been noted, although whether these translate to altered outer membrane permeability or vesicle production remains to be determined. The rapid growth rate of PCC 11901, linked to lower photoinhibition and higher photosynthetic rates[11–13], may also contribute through increased membrane dynamics during cell division, as faster-dividing cells undergo more frequent rounds of membrane remodelling that could facilitate outer membrane transit of periplasmic proteins.

The clear superiority of FutA over all five Sec-pathway candidates tested for eYFP secretion is likely multifactorial. One probable factor is the capacity of the Tat pathway to translocate fully folded proteins, enabling cytoplasmic chromophore maturation in eYFP before membrane translocation. Sec-mediated export threads unfolded polypeptides through the SecYEG translocon, exposing them to the oxidising periplasmic environment where aberrant disulfide bond formation can interfere with β-barrel assembly and chromophore cyclisation [32,33]. Because our assay measured fluorescence rather than total protein, the relative contributions of transport efficiency and post-translocation folding fidelity cannot be deconvolved from the present dataset. Importantly, however, the poor performance of PhoX (17.1%), the second Tat candidate tested, demonstrates that Tat pathway access alone is insufficient for efficient secretion, and that signal peptide-specific features play a dominant role. Analysis of the two Tat signal peptides offers insight into why FutA vastly outperformed PhoX. Although both carry a net charge of +5 and a canonical twin-arginine motif, they differ markedly in architecture. The FutA signal peptide (32 residues, GRAVY +0.27) has a short N-region of only 8 residues before the RR motif, followed by a well-defined 16-residue hydrophobic core (FFVGGTALTALVVANL). The PhoX signal peptide (57 residues, GRAVY −0.30) is nearly twice as long, with an extended 27-residue N-region containing three negatively charged residues (Asp/Glu) before the RR motif, and an overall hydrophilic character. This extended, polar N-region may reduce the efficiency of TatBC receptor recognition and membrane targeting, consistent with the established requirement for a compact, positively charged N-region and a sufficiently hydrophobic H-region for efficient Tat translocation. The cargo-dependent reversal observed in Phase 2, where thermitase outperformed FutA for lichenase secretion by 2.6-fold, can be rationalised by considering the biophysical properties of the two target proteins. eYFP (26.9 kDa, pI 5.6) is a rigid 11-stranded β-barrel that requires cytoplasmic folding and autocatalytic chromophore maturation before it becomes fluorescent; its two cysteine residues are also susceptible to aberrant disulfide bond formation in the oxidising periplasmic environment [32,33]. These properties strongly favour Tat-mediated export, which translocates the fully folded, fluorescence-competent protein. Lichenase from *C. thermocellum* (∼36 kDa, 334 residues, pI 4.8) is, by contrast, a thermostable (β/α)_8_ TIM barrel enzyme with an activity optimum at 65 °C that does not require cofactors or post-translational modifications for enzymatic function. Its inherent thermostability likely enables rapid and efficient refolding after Sec-mediated translocation in an unfolded state, explaining why the Sec-pathway leader thermitase outperformed FutA for this particular cargo. Together, these observations underscore that no single signal peptide is universally optimal: Tat leaders may be favoured for folding-dependent cargo proteins, while Sec leaders may be preferable for thermostable enzymes that refold efficiently post-translocation.

The growth inhibition observed in signal peptide-bearing strains was comparable between Tat and Sec constructs (Section 3.3), indicating that the metabolic cost of Tat-mediated export is not prohibitively higher than Sec-mediated secretion under the conditions tested. This is an encouraging finding for industrial applications, as it suggests that the superior secretion efficiency of FutA does not come at a disproportionate cost to biomass accumulation.

Among the Sec candidates, the superior performance of the heterologous thermitase signal peptide over all four native PCC 11901 leaders is noteworthy and consistent with the observation by Nandru et al. (2025) [27] that thermitase drives effective eYFP secretion in *C. aponinum* PCC 10605 using a similar RSF1010-based expression system [27]. This result suggests that heterologous signal peptides should be routinely included in future secretion engineering campaigns in PCC 11901. One plausible explanation is that native signal peptides have co-evolved with their endogenous substrates and may exhibit suboptimal interactions with heterologous proteins of different size, charge, or folding kinetics. Empirical screening across both native and heterologous panels therefore remains essential.

The extension of the platform from eYFP (∼27 kDa) to lichenase (∼36 kDa) demonstrates that the system accommodates functionally diverse protein classes. Quantitative assessment of extracellular lichenase activity using the K-MBG4 chromogenic assay revealed activities of approximately 74.99 ± 1.78 mU/mL/OD_730_ for thermitase and 28.83 ± 0.96 mU/mL/OD_730_ for FutA. To our knowledge, these represent the first quantitative measurements of secreted enzymatic activity in any cyanobacterium. The only prior report of lichenase secretion in cyanobacteria, by Sergeyenko *et al*. (2003) [26] in *Synechocystis* sp. PCC 6803, relied exclusively on qualitative Congo Red plate assays without reporting activity units [26], and the only other quantitative secretion yield reported for a cyanobacterium is the 779 ± 40 µg L^−1^ estimated by densitometry for a lytic polysaccharide monooxygenase in *S. elongatus* UTEX 2973 [22]. The present work therefore establishes the first quantitative enzymatic benchmark for cyanobacterial heterologous protein secretion.

While the absolute activities remain below those achievable in optimised heterotrophic hosts, for example, secretion of a chimeric *C. thermocellum* lichenase from *Bacillus subtilis* has been reported at 80.56 U/mL through signal peptide hydrophobicity engineering [34], this comparison must be considered in context, acknowledging differences in cultivation regimes, normalisation metrics, and the fundamentally distinct envelope architectures of Gram-positive and diderm organisms. PCC 11901 is a diderm cyanobacterium in which secreted proteins must traverse both the cytoplasmic and outer membranes, whereas *B. subtilis* possesses a single membrane and has benefitted from decades of secretion optimisation. Moreover, PCC 11901 offers the distinct advantage of photoautotrophic production independent of organic carbon feedstocks, and the activities detected here validate the secretion platform as a functional system amenable to further optimisation through promoter engineering, signal peptide combinatorics, and culture condition development.

Notably, the rank order of signal peptide performance was reversed relative to Phase 1: whereas FutA substantially outperformed thermitase for eYFP secretion (92.2% vs 55.7%), thermitase directed approximately 2.6-fold higher extracellular lichenase activity than FutA. This cargo-dependent reversal is consistent with the distinct translocation mechanisms of the two pathways: the Tat pathway preferentially exports folded substrates such as eYFP, whereas the Sec pathway may more efficiently handle proteins like lichenase that fold rapidly post-translocation and do not require cytoplasmic maturation for enzymatic activity. The lichenase experiments permit a direct comparison with Sergeyenko *et al*. (2003) [26], who first demonstrated heterologous protein secretion in cyanobacteria using the same enzyme from *C. thermocellum* in *Synechocystis* sp. PCC 6803. Our work extends those findings in three important respects. First, we demonstrate lichenase secretion in the faster-growing PCC 11901, which offers superior biomass accumulation and industrial scalability. Second, we employ signal peptides that engage the canonical Sec (thermitase) and Tat (FutA) translocation pathways rather than pilin-type leaders whose routing through the Type IV pilus system is now considered distinct from general protein secretion [23,24]. Third, we provide quantitative activity measurements using the K-MBG4 chromogenic assay in addition to the qualitative Congo Red assay used by Sergeyenko *et al*. (2003) [26].

A question left open by these experiments concerns outer membrane transit. The Sec and Tat pathways translocate proteins across the cytoplasmic membrane, but the mechanism by which proteins subsequently traverse the outer membrane remains uncharacterised. Several non-exclusive possibilities include outer membrane vesicle (OMV) release [35], passage through porins, or unidentified secretion apparatus. Notably, our fluorescence-based assay does not distinguish between soluble extracellular eYFP and eYFP associated with extracellular vesicles (EVs). Recent work in *Synechocystis* sp. PCC 6803 has demonstrated that periplasmic proteins can reach the extracellular space both as soluble protein and encapsulated within EVs [35]. Determining the relative contribution of vesicle-mediated versus soluble export in PCC 11901, for example by differential ultracentrifugation of culture supernatants, would be an important next step to fully characterise the secretion mechanism. Recent exoproteomic data indicate that the T4P system is dedicated to pilin secretion rather than serving as a general conduit [24], and quantitative secretion assays in *Synechocystis* confirmed that NanoLuc reporter secretion is Independent of T4P assembly [23]. The successful extracellular recovery of two proteins of different sizes and properties suggests that the PCC 11901 outer membrane is more permissive than that of PCC 6803, consistent with the architectural differences revealed by comparative genomics [11].

### Limitations and considerations

Several aspects of the experimental design merit discussion. First, all constructs were expressed from the self-replicating RSF1010 plasmid. To determine whether chromosomal integration would improve expression, we compared eYFP fluorescence from RSF1010 versus integration at the validated *mrr* neutral site [13] under the constitutive P*cpc560*. RSF1010-based expression yielded approximately 2.8-fold higher fluorescence per OD_730_ than chromosomal integration at 24 hours (; **Supplementary Figure S10**). These results contrast with a previous report of higher expression from chromosomal loci in PCC 11901 [13], a discrepancy that may reflect differences in growth conditions or locus-specific effects at the *mrr* site. Critically, this finding demonstrates that the secretion efficiencies reported here could not be improved simply by moving to chromosomal integration, and that RSF1010 represents a validated expression platform for this strain. The copy number of RSF1010 in PCC 11901 has not been determined. The chromosome copy number, however, has recently been estimated at approximately 4-7 in wild-type cells by flow cytometry and shown to vary with growth phase, being highest in early exponential phase and declining thereafter [14]. This polyploidy implies that chromosomally integrated constructs are present at multiple copies per cell, yet RSF1010 still outperformed genomic integration, suggesting that plasmid copy number exceeds the chromosome count under the conditions tested. The dynamic nature of chromosome copy number may also contribute to the inter-replicate variability observed across time points. Second, the non-secreting control (eYFP without signal peptide) did exhibit low-level extracellular fluorescence (9.3 ± 2.9% of total fluorescence at day 7), indicating that a degree of cell lysis occurs under the culture conditions employed. However, this baseline was substantially lower than the extracellular fluorescence observed for any signal peptide construct. The signal peptide-dependent rank order of secretion efficiencies, with FutA at 92.2% and several Sec candidates below 20%, is inconsistent with a lysis-driven mechanism, which would produce comparable extracellular fluorescence across all strains regardless of signal peptide identity. Furthermore, the temporal profile of extracellular fluorescence in the non-secreting control peaked at day 2 and declined thereafter, mirroring the intracellular fluorescence trajectory, whereas FutA-driven extracellular fluorescence increased continuously over 7 days, a pattern indicative of active export rather than passive release. Third, extracellular proteases could degrade secreted proteins in the culture medium, potentially leading to underestimation of true secretion efficiency. Fourth, as noted in Section 3.4, the unequal representation of Sec (n = 5) and Tat (n = 2) signal peptides means that the present data establish FutA as the best individual signal peptide rather than demonstrating broad pathway-level superiority of Tat over Sec.

From a translational perspective, extracellular secretion eliminates the requirement for cell disruption, reduces purification steps, and enables continuous product harvesting. The thermodynamic case for cyanobacterial biomanufacturing has been further strengthened by recent analyses demonstrating that *Synechococcus* sp. biomass formation captures ∼275 kJ per mol of CO_2_ fixed as an enthalpy change, energy that is theoretically recoverable in the secreted product [9]. Combined with the exceptional growth kinetics of PCC 11901 (doubling time ∼2 h), its halotolerance, its capacity for high cell-density cultivation to >33 g DCW L^−1^ [10,11], and the comprehensive genetic engineering toolbox now available [13], these results position PCC 11901 as one of the most promising cyanobacterial chassis for scalable biomanufacturing of extracellular recombinant proteins.

### Future directions

Several avenues could further enhance the utility of this platform. Although our comparative data indicate that RSF1010-based expression currently outperforms *mrr* integration for the constructs tested here (**Supplementary Figure S10**,**S11**), the superiority of the self-replicating plasmid over chromosomal integration was not restricted to constitutive expression. Under a separate IPTG-inducible promoter system, the self-replicating construct induced with 100 µM IPTG yielded approximately 4-fold higher eYFP fluorescence than the equivalent integrative construct (**Supplementary Figure S11**), confirming that the plasmid-based platform outperforms *mrr* integration across both constitutive and inducible regulatory architectures. Evaluation of alternative neutral sites such as *aquI* or *glgA1* [13] may yield different results and would eliminate plasmid maintenance costs and the requirement for antibiotic selection. The tightly regulated DAPG-inducible *PhlF/PphlF* promoter system characterized in PCC 11901 [13], could enable temporal control of secretion, decoupling biomass accumulation from product export. Expansion of the Tat signal peptide panel, guided by bioinformatic mining of PCC 11901’s 3,316 predicted proteins, could identify native Tat leaders that match or exceed the performance of FutA. Complementary mass spectrometry-based analysis of signal peptide cleavage sites would confirm correct processing and enable optimization of the signal peptide-target protein junction. A combinatorial approach in which FutA is used as the default for folding-dependent proteins and thermitase for intrinsically disordered or rapidly folding targets could maximise the range of secretable products. Differential ultracentrifugation or size-exclusion chromatography of culture supernatants could determine whether the secreted proteins are released as soluble molecules or associated with extracellular vesicles, a distinction with implications for both mechanistic understanding and downstream purification strategies. Future work should also include quantification of extracellular lichenase protein by Western blot or ELISA to separate secretion yield from post-translocation functional integrity, enabling a direct comparison of secretion efficiency across cargo proteins. Finally, scaling of PCC 11901 secretion cultures in photobioreactors, for which photo-mechanistic models are already available [12], would provide the data needed to assess industrial feasibility.

## 5. Conclusions

We have established *Synechococcus* sp. PCC 11901 as a tuneable platform for recombinant protein secretion through a systematic two-phase experimental programme. Phase 1 identified FutA (Tat pathway) and thermitase (Sec pathway) as the most effective signal peptides from a panel of seven candidates, with FutA achieving the highest secretion efficiency for eYFP (92.2 ± 0.3% extracellular export by day 7). Daily profiling over seven days revealed distinct kinetic signatures: FutA-driven secretion increased continuously without saturation, whereas Sec-mediated export plateaued by day 3. Phase 2 confirmed that the platform supports enzymatic target proteins, with both FutA and thermitase directing active lichenase to the extracellular fraction. Notably, thermitase outperformed FutA for lichenase secretion by 2.6-fold, demonstrating that optimal signal peptide selection is cargo dependent. Together, these results provide a rational framework for engineering extracellular protein production in cyanobacteria and demonstrate that signal peptide selection with the Tat-pathway leader FutA as the preferred default for folding-dependent cargo and thermitase for Sec-compatible targets, constitutes a powerful, transferable strategy for secretion optimization.

## Supporting information

Supplementary Figure S1.Intracellular eYFP fluorescence time course

Supplementary Figure S2. Whole-culture eYFP fluorescence time course

Supplementary Figure S3. Chemidoc UV Images futA fluorescence time course

Supplementary Figure S4.% secreted Day 7 bar chart

Supplementary Figure S5. Extracellular Day 7 bar chart

Supplementary Figure S6.Grouped daily extracellular eYFP fluorescence

Supplementary Figure S7.Grouped daily secretion efficiency for all constructs

Supplementary Figure S8. Congo Red colony overlay assay

Supplementary Figure S9.MBG4 raw absorbance at 400 nm

Supplementary Figure S10. RSF1010 vs mrr under constitutive expression (Pcpc560, 24h)

Supplementary Figure S11. RSF1010 vs mrr under IPTG-inducible expression

Supplementary Table S1.Signal peptide sequences and properties

## Supplementary Materials

The following supporting information can be downloaded at: [link]. Figure S1: Intracellular eYFP fluorescence time course; Figure S2: Whole-culture eYFP fluorescence time course; Figure S3: ChemiDoc UV images of FutA extracellular fractions days 1–7; Figure S4: Secretion efficiency (% secreted) at day 7; Figure S5: Extracellular eYFP fluorescence at day 7; Figure S6: Grouped daily extracellular eYFP fluorescence; Figure S7: Grouped daily secretion efficiency for all constructs; Figure S8: Congo Red colony overlay assay; Figure S9: MBG4 raw absorbance at 400 nm; Figure S10: RSF1010 vs mrr integration under constitutive expression; Figure S11: RSF1010 vs mrr integration under IPTG-inducible expression; Table S1: Signal peptide sequences and properties.

## Author Contributions

Conceptualization, J.Á.M.-C.; methodology, J.Á.M.-C. and A.B.; validation, J.Á.M.-C. and A.B.; formal analysis, J.Á.M.-C.; investigation, J.Á.M.-C. and A.B.; resources, U.S.S.; data curation, J.Á.M.-C.; writing—original draft preparation, J.Á.M.-C. and A.B.; writing—review and editing, J.Á.M.-C., A.B., D.L., K.G., D.S.K. and U.S.S.; visualization, J.Á.M.-C. and A.B.; supervision, J.Á.M.-C. and U.S.S.; project administration, J.Á.M.-C. and U.S.S.; funding acquisition, U.S.S. All authors have read and agreed to the published version of the manuscript.

## Institutional Review Board Statement

Not applicable.

## Informed Consent Statement

Not applicable.

## Data Availability Statement

The data that support the findings of this study are available from the corresponding author upon reasonable request.

## Acknowledgments

The authors thank Dr David Lea-Smith (University of East Anglia) for critical review of the manuscript. During the preparation of this manuscript, the authors used AI-assisted writing tools for the purposes of language editing. The authors have reviewed and edited the output and take full responsibility for the content of this publication.

## Disclaimer/Publisher’s Note

The statements, opinions and data contained in all publications are solely those of the individual author(s) and contributor(s) and not of MDPI and/or the editor(s). MDPI and/or the editor(s) disclaim responsibility for any injury to people or property resulting from any ideas, methods, instructions or products referred to in the content.

