## Supplementary figures and images for "Systematic Evaluation of Signal Peptide-Driven Protein Secretion in the Fast-Growing Cyanobacterium *Synechococcus* sp. PCC 11901"

### Supplementary Figure S1.Intracellular eYFP fluorescence time course

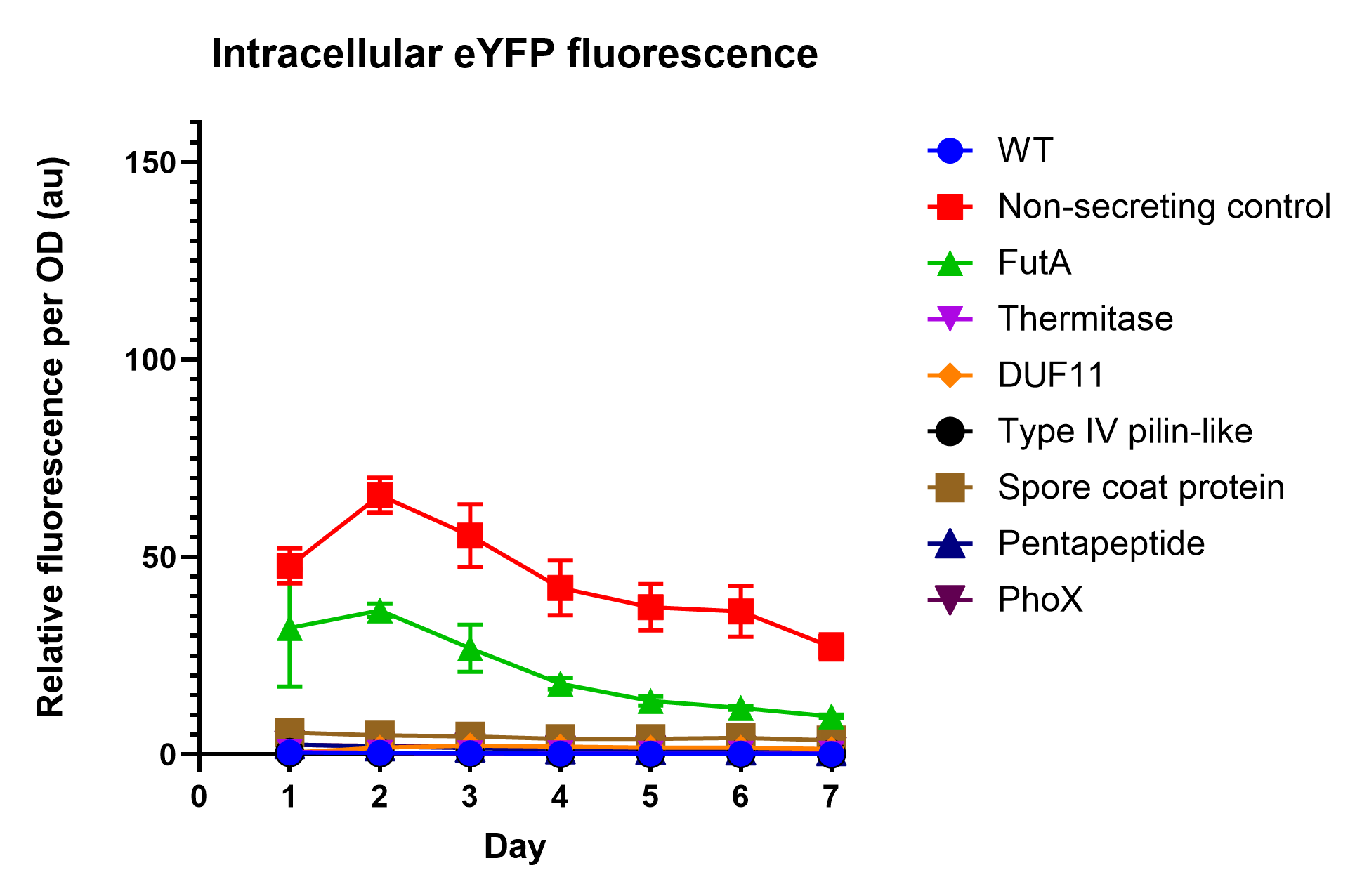

### Supplementary Figure S2. Whole-culture eYFP fluorescence time course

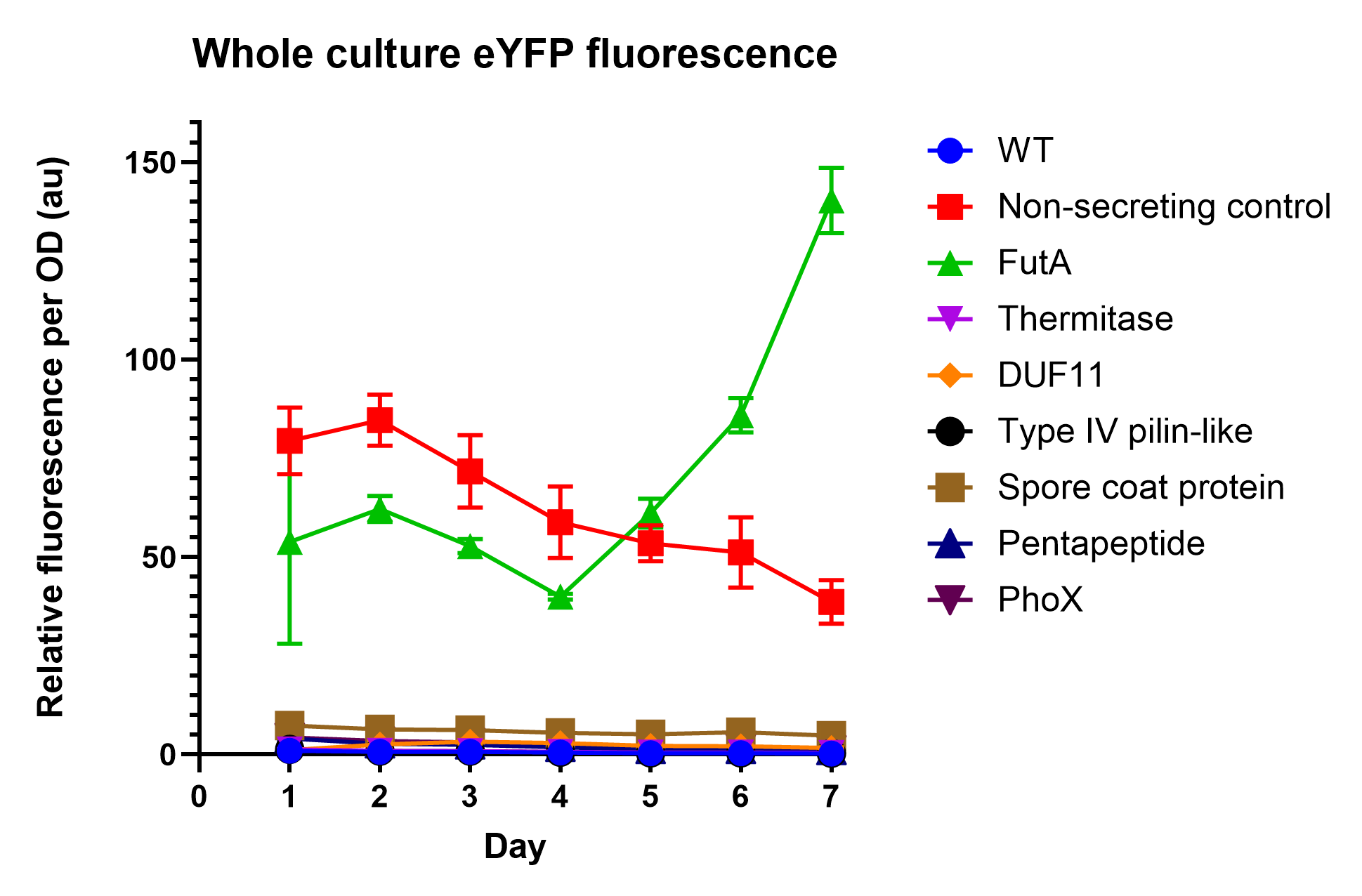

### Supplementary Figure S3. Chemidoc UV Images futA fluorescence time course

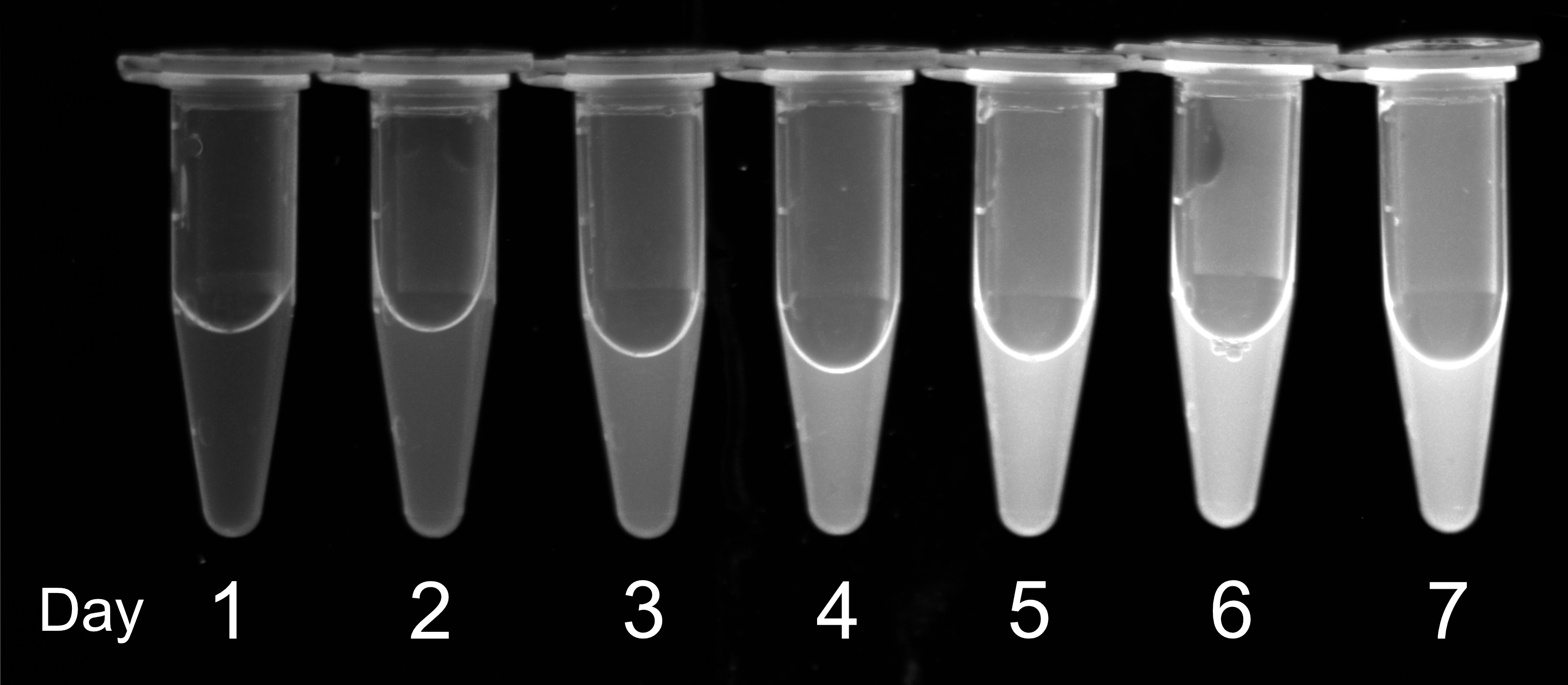

### Supplementary Figure S4.% secreted Day 7 bar chart

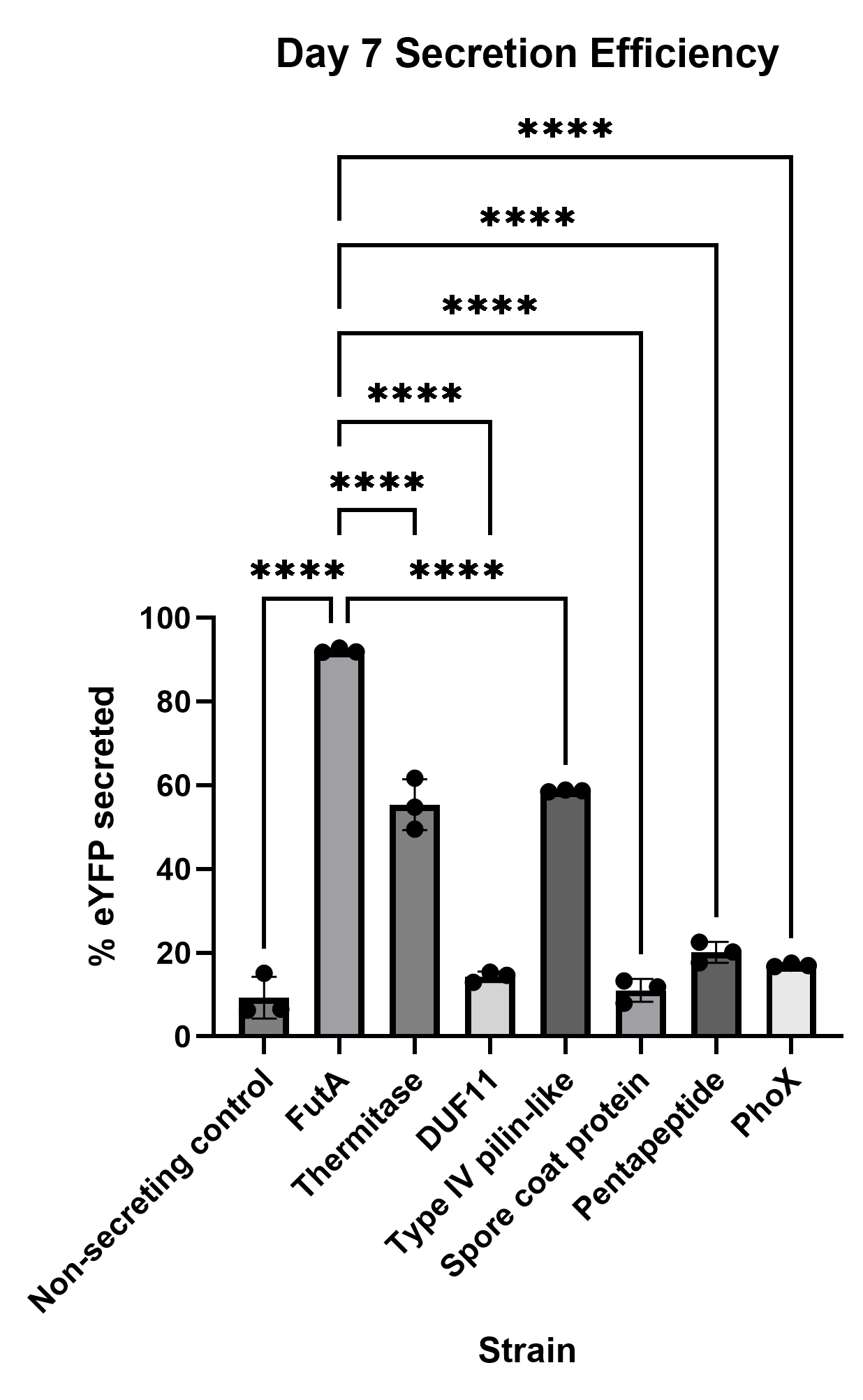

### Supplementary Figure S5. Extracellular Day 7 bar chart

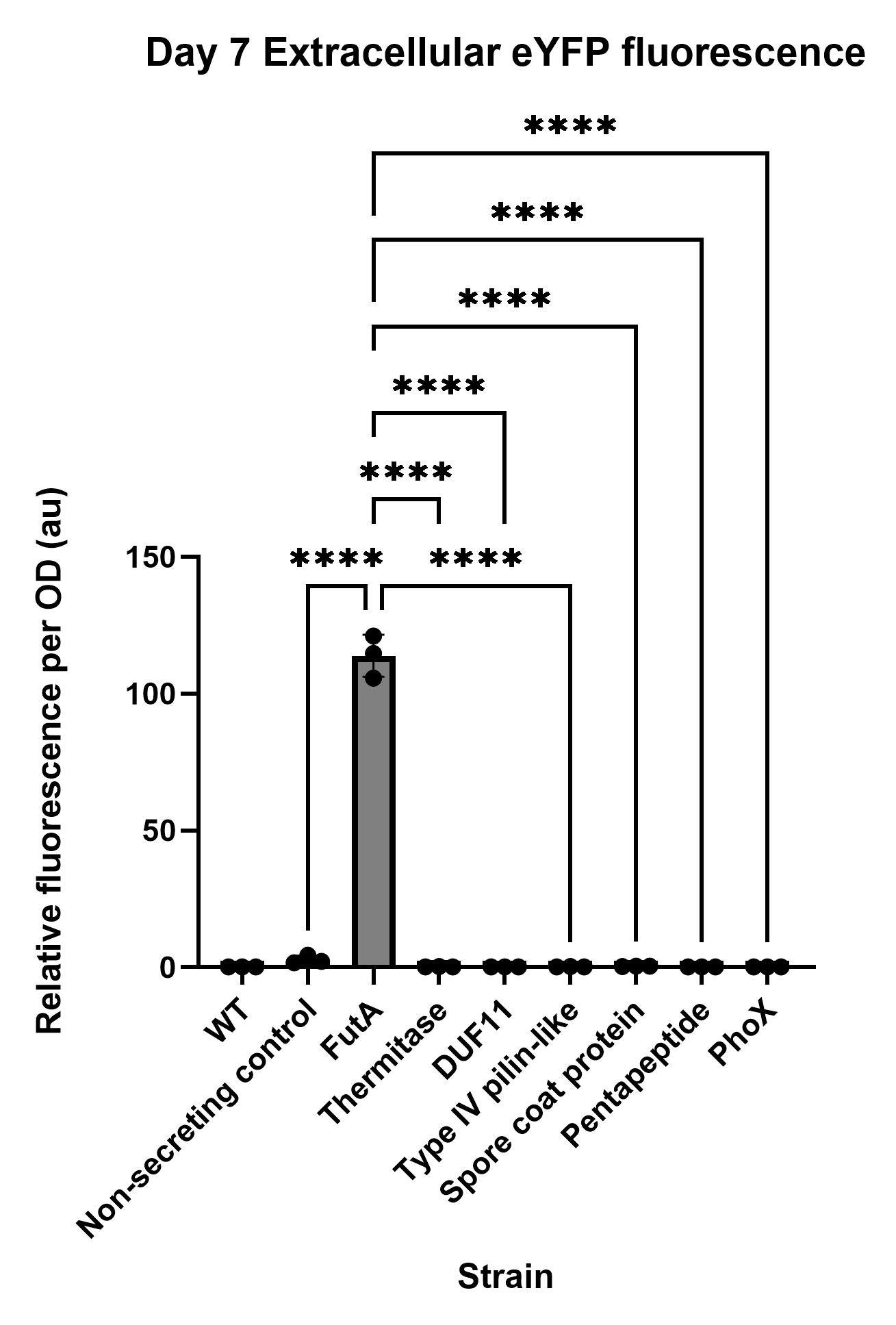

### Supplementary Figure S6.Grouped daily extracellular eYFP fluorescence

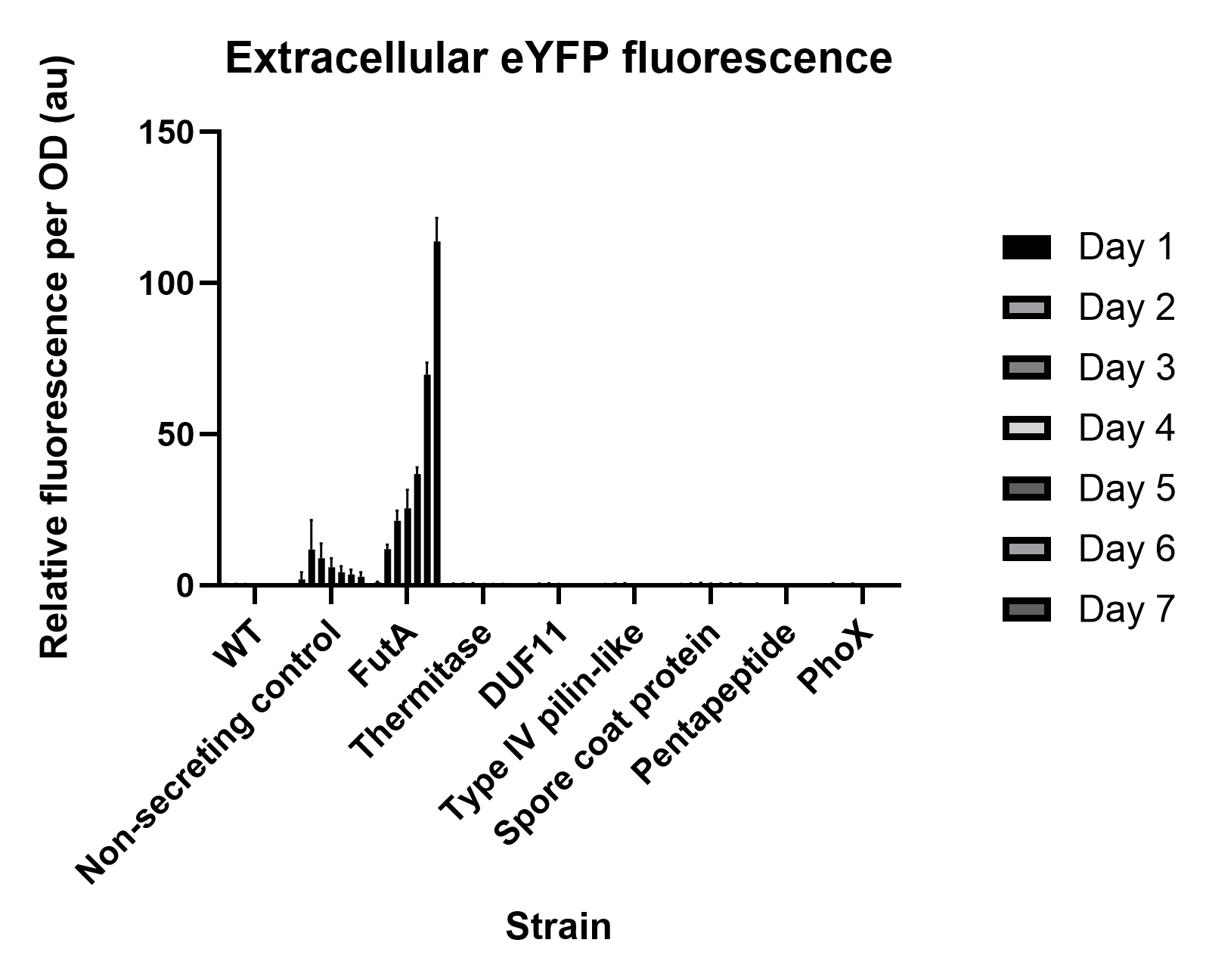

### Supplementary Figure S7.Grouped daily secretion efficiency for all constructs

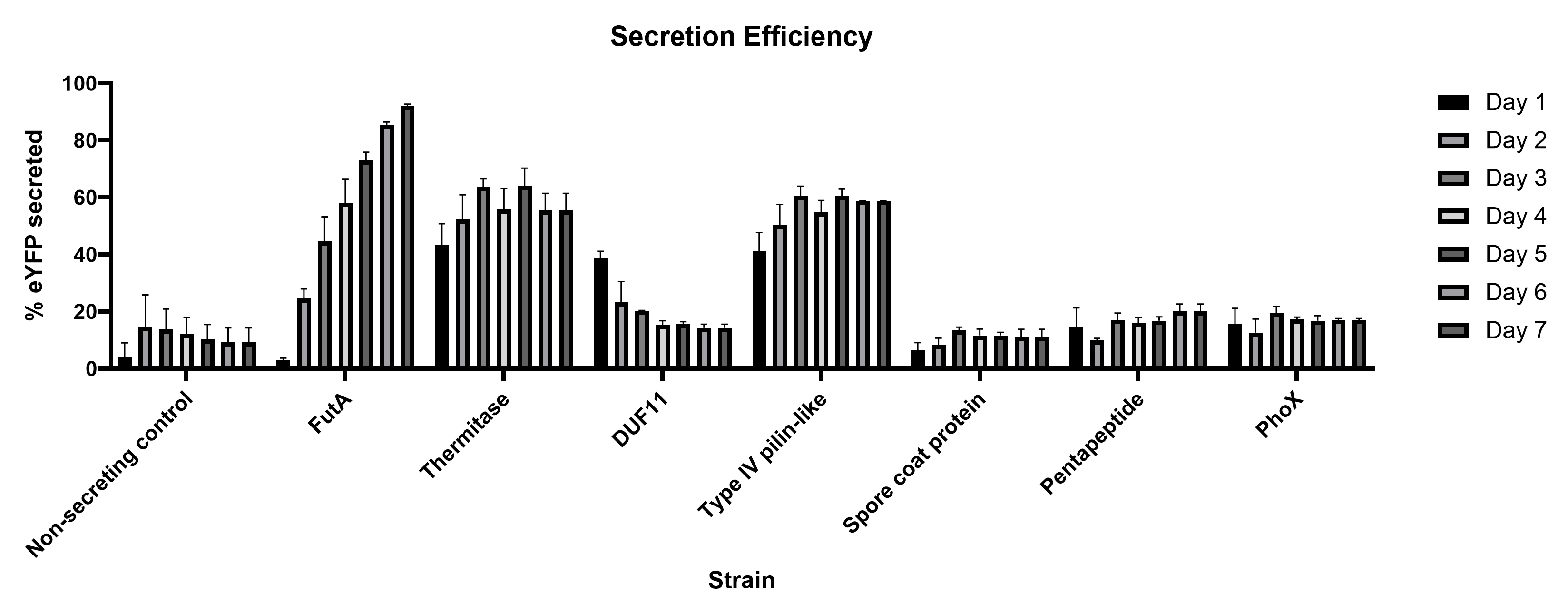

### Supplementary Figure S8. Congo Red colony overlay assay

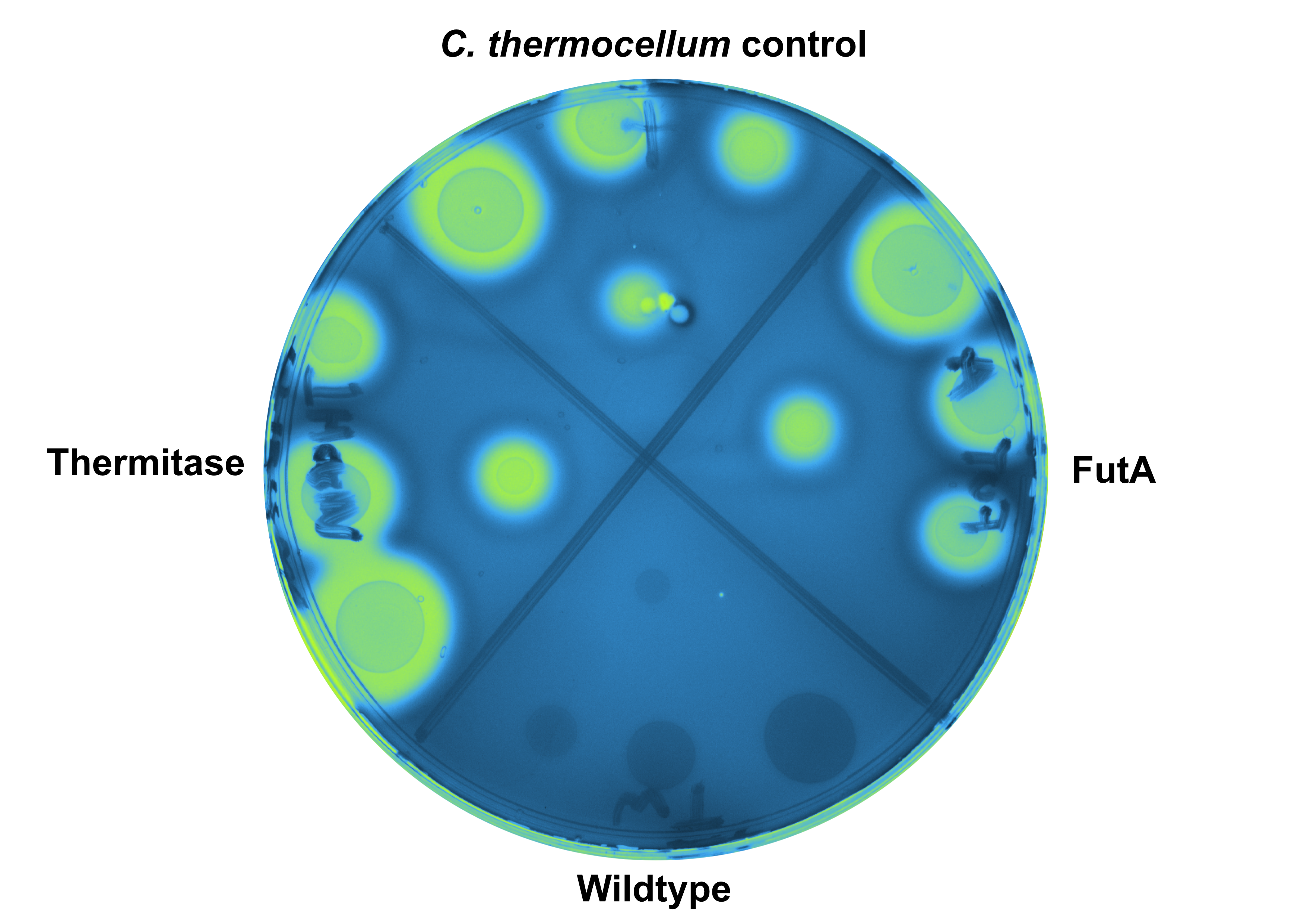

### Supplementary Figure S9.MBG4 raw absorbance at 400 nm

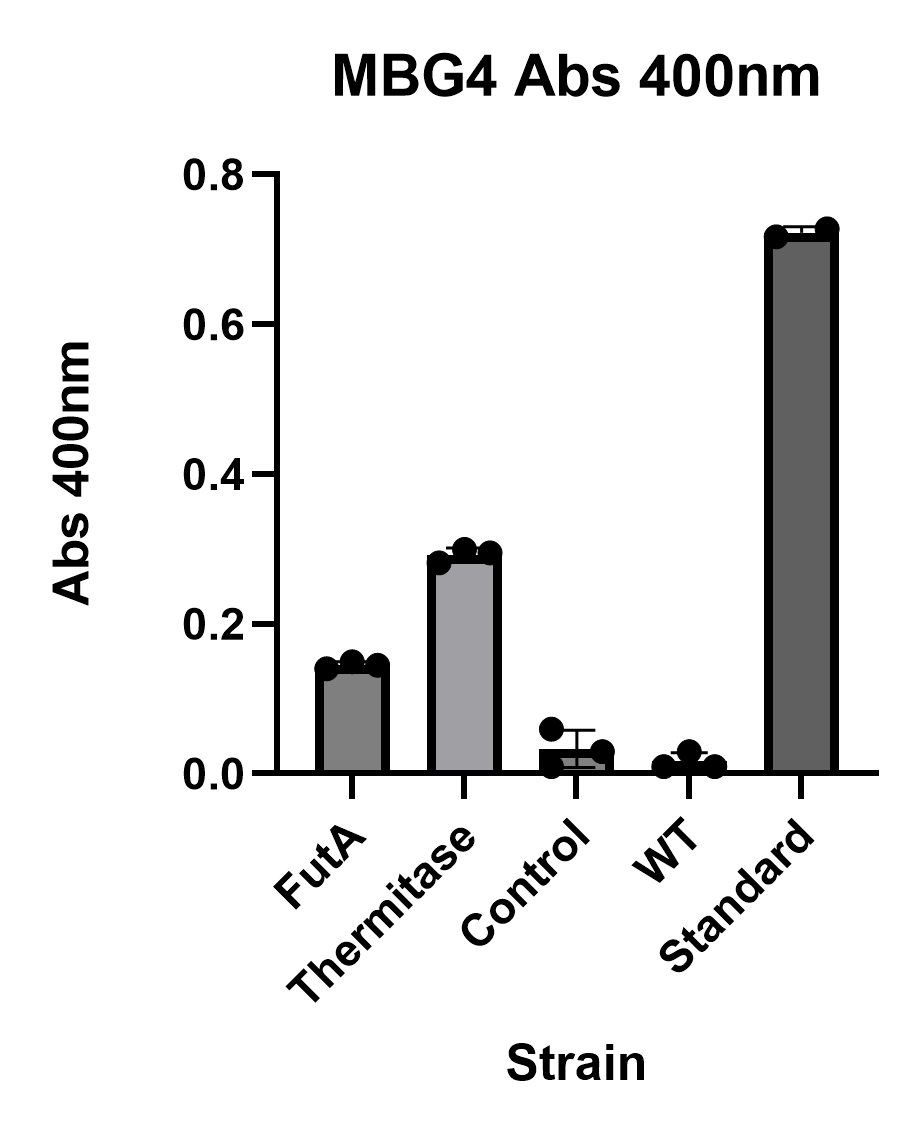

### Supplementary Figure S10. RSF1010 vs mrr under constitutive expression (Pcpc560, 24h)

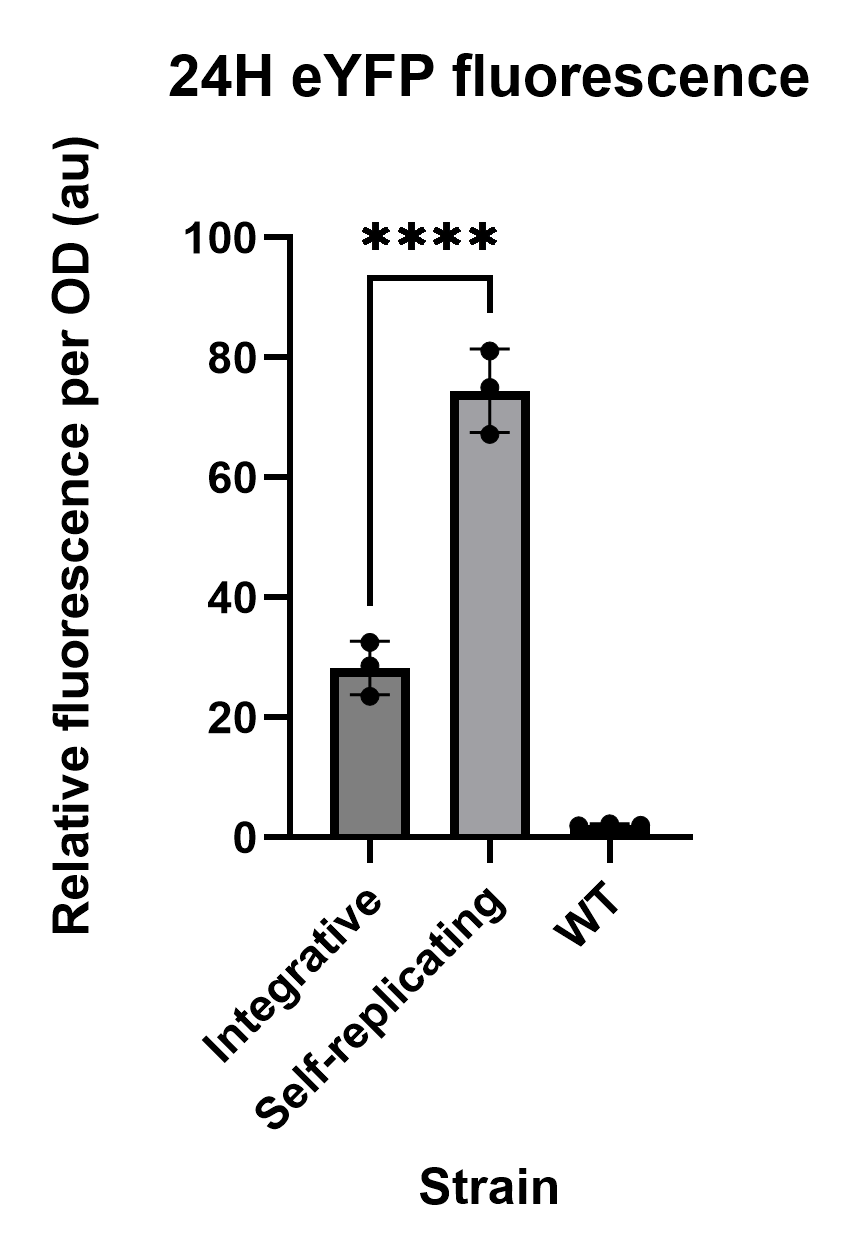

### Supplementary Figure S11. RSF1010 vs mrr under IPTG-inducible expression

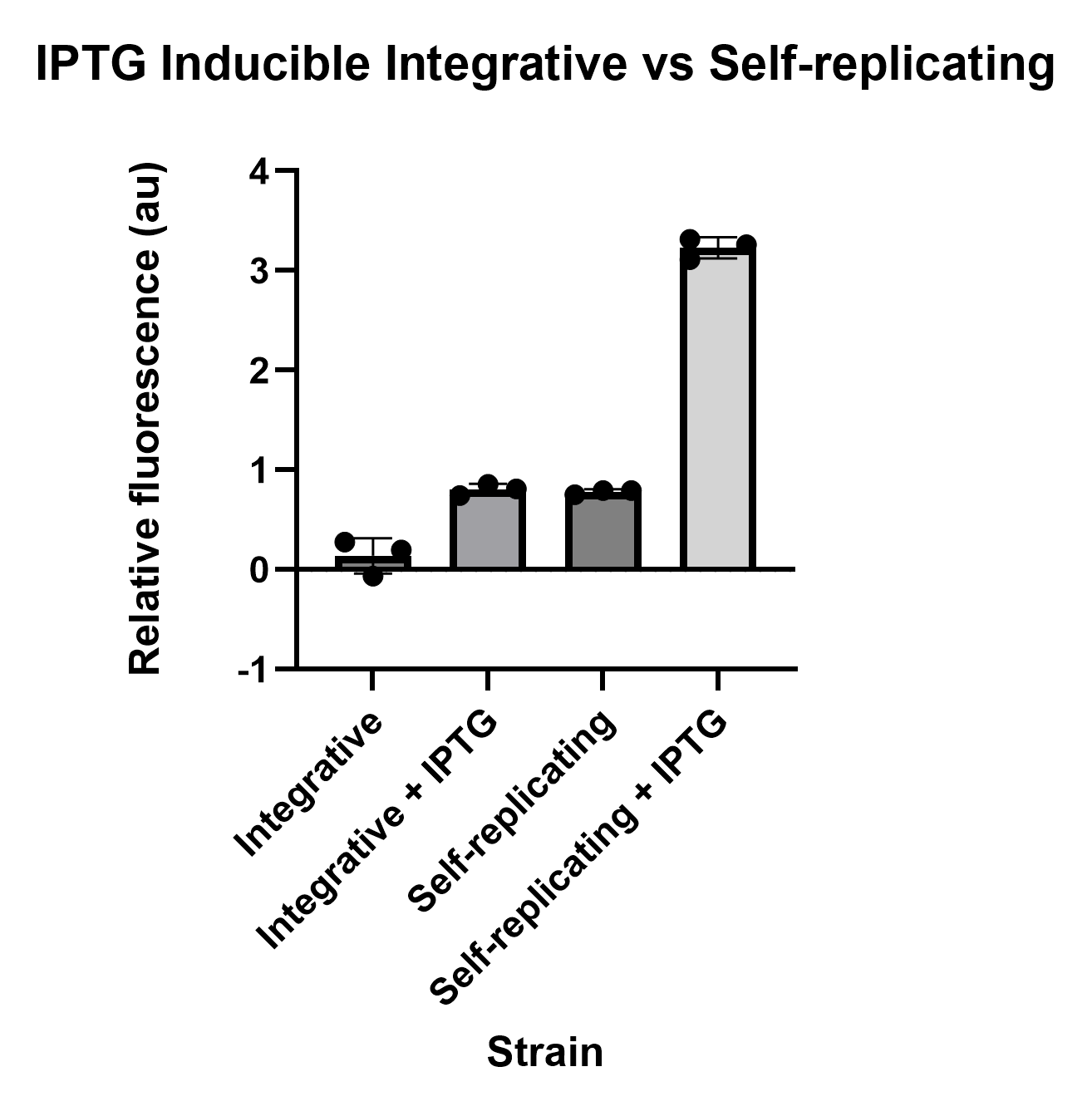
